# A dynamical low-rank approach to solve the chemical master equation for biological reaction networks

**DOI:** 10.1101/2022.05.04.490585

**Authors:** Martina Prugger, Lukas Einkemmer, Carlos F. Lopez

## Abstract

Solving the chemical master equation is an indispensable tool in understanding the behavior of biological and chemical systems. In particular, it is increasingly recognized that commonly used ODE models are not able to capture the stochastic nature of many cellular processes. Solving the chemical master equation directly, however, suffers from the curse of dimensionality. That is, both memory and computational effort scale exponentially in the number of species. In this paper we propose a dynamical low-rank approach that enables the simulation of large biological networks. The approach is guided by partitioning the network into biological relevant subsets and thus avoids the use of single species basis functions that are known to give inaccurate results for biological systems. We use the proposed method to gain insight into the nature of asynchronous vs. synchronous updating in Boolean models and successfully simulate a 41 species apoptosis model on a standard desktop workstation.

## 1. Introduction

Mechanistic models are commonly used to acquire insights about the biochemical reaction networks that govern cellular processes inside a cell. These models are typically obtained by a combination of expert and literature-driven knowledge as well as experimental data. Models based on ordinary differential equations (ODE) are most commonly used [12]. However, it is also well known that in a number of problems the ODE formulation is insufficient to describe important features of the biological system [48, 25, 36, 38]. Often this inability comes from the fact that ODE models replace stochastic dynamics by some average. In contrast, such stochastic effects are taken into account by models based on the chemical master equation (CME).

The primary challenge associated with models based on the CME is the large computational cost. For a direct solver, the computational effort and memory requirements increase exponentially with the number of species, limiting such simulations to relatively small systems. This is a reason why the less computationally expensive ODE models are popular. In this paper, we employ a dynamical low-rank approximation to drastically reduce the computational and memory costs and therefore make the solution of large, complex systems tractable. In our approach, the network is divided into subsets that are tightly coupled, while only averaged information is exchanged between them. This enables simulations with a network size that would be prohibitive for direct solvers. Although low-rank approximations have been used in the past - particularly in the quantum physics literature - our approach departs from traditional applications by exploiting the underlying structure of the network rather than reducing the equations to single species basis functions (orbitals in the language of quantum dynamics). This represents a significant shift in how these methods should be applied in order to obtain accurate results with drastically reduced computational effort.

We note that in the literature the large computational cost of solving the CME has been addressed in a variety of ways. If a system is very sparse (i.e. only a few chemical states are populated) finite state projection 35 is a viable way to reduce computational complexity. However, especially for Boolean models (for details see below) this essentially assumes that a large number of species is either on or off with probability 1, a very unrealistic assumption. The other commonly used technique is Monte-Carlo simulation or SSA [22, 26]. In this approach trajectories in the space of possible configurations are sampled and a statistical representation is obtained by repeated simulations. Thus, to obtain, e.g., low abundant phenotypes requires the computation of an excessively large number (depending on the problem potentially tens to hundreds of thousands) of trajectories [23].

The idea of dynamical low-rank approximation is to use lower dimensional basis functions in order to approximate the high dimensional probability distribution. If a relatively small number of such basis functions is sufficient to describe the relevant dynamics, this results in a drastic reduction of the required computational resources. In Jahnke et al. [27], such an approach is presented for a relatively low-dimensional CME. In this work, following the quantum physics literature, each species is considered its own subspace (i.e. the basis functions only depend on a single species). The proposed method works relatively well for the given examples (*≈* five species and *≈* ten reactions), but the authors are somewhat discouraging regarding the wider use of the method. The reason for this is that in biological applications, it would be difficult or even impossible to consider each species as independent, particularly given the intricate structures of complex biological networks 3. Therefore, approximating the probability distribution of a system of reactions using single species basis function could significantly degrade the accuracy of the method. Our approach avoids this by using insight into the structure of the reaction network when choosing the dependence of the basis functions. This results in a dynamical low-rank algorithm that can accurately simulate many real world reaction networks at low computational cost.

Our proposed dynamical low-rank approximation formalism for biological networks leverages prior work. In particular, the projector splitting based dynamical low-rank approach introduced by Lubich et al. [29] and its later extension to hierarchical tensors [30]. This was the first robust low-rank approach able to handle dynamical systems [34]. More recently, this approach found success in solving particle kinetic equations such as the Vlasov equation [18, 19, 17] and radiative transfer problems [40, 15, 14, 16]. Our dynamical low-rank approach is fully deterministic (i.e. noise free) and provides the entire probability distribution with a single simulation. This enables us to get an accurate picture of the network dynamics, while still being relatively cheap from a computational point of view. In addition, the proposed approach does not show unfavorable scaling with the number of reactions (as does SSA), but only with the dependent species for each reaction. The approximation error in the dynamical low-rank approach is controlled by how the network is partitioned, thus allowing us to make use of expert knowledge to refine the result.

While the dynamical low-rank approach can be applied to any model based on the CME, in this paper we consider Boolean models [54]. In this formalism, each biochemical species (e.g. genes, proteins) can exist in two states (on or off), which significantly simplifies model complexity and only requires that one knows the structure of the network (which in some situations can be inferred from data 41). The state of the network is then determined by a binary string that encodes which species are switched on and which are switched off. In the CME approach a probability is assigned to each such state. Boolean models have proven very useful to extract mechanistic insights from complex systems 52, 13 and a range of software packages are available for practitioners. We note that the Boolean model formalism is also used in a variety of applications outside biology including quantum physics, circuit theory, natural language processing, and search engines 37, 5. For increasingly large and complex problems Boolean models have the added advantage that they do not require precise knowledge of a large number of kinetic parameters, which otherwise must be either measured experimentally, or inferred using statistical methods 46, 2.

## 2. Dynamical low-rank approximation

### 2.1. Setup

The chemical master equation is given by

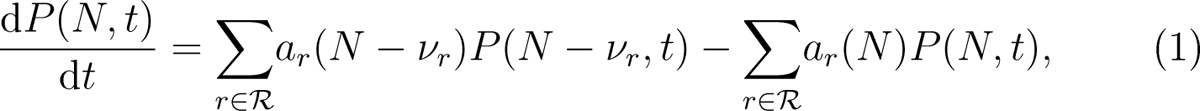

where *N* (*t*) = (*N*_1_(*t*)*, N*_2_(*t*)*, …, N_n_*(*t*)) and *N_i_* is the number of molecules of the chemical species of type *i*. The number of distinct chemical species is denoted by *n*. *P* (*N, t*) describes the state of the system determined by the set of all possible chemical reactions *R* and the reaction propensities *a_r_*(*N*), where *ν_r_*represents the change of species due to reaction *r ∈ R*.

In the case of a Boolean model, instead of number of molecules, we have states *x_i_ ∈ {*0, 1*}*, where each *x_i_* represents the state of a chemical species in the system. That is, each chemical species can only be in one of two states (often referred to as on and off, or active and inactive). The chemical master equation describes the dynamics (i.e. time evolution) of a probability distribution *P* (*t, x*_1_*, …, x_d_*). Boolean models are often specified by a set of (logic) rules. For an explanation of the translation of a Boolean ruleset describing a biological network into a system of ODEs, we refer the reader to Appendix A.

To apply the dynamical low-rank algorithm we partition the *d* nodes of the network into two groups containing *{x*_1_*, …, x_m_}* and *{x_m_*_+1_*, …, x_d_}* respectively. Ideally, the two partitions are as equal as possible (i.e., *m* = *d/*2).

Note, that within this framework we can use an arbitrary partition of the network by simply renaming the nodes. The performance of the algorithm is crucially dependent on how this partitioning is done. Both domain specific knowledge (i.e. knowledge on the behavior of the reaction network) and mathematical techniques (such as graph partitioning algorithms) can be used to find good partitionings. We will discuss this in more detail later in the paper.

We now write *P* in matrix form by linearizing the elements within each partition. That is, we linearize a state according to the function.

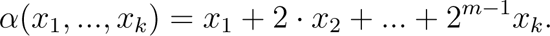

We then consider the following matrix

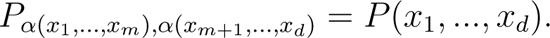

This means, that the index of the state in the first partition determines the row index in the matrix, and the index of the state in the second partition determines the column index.

To address the curse of dimensionality, we avoid forming the full matrix *P* (i.e. nowhere in the algorithm will it be necessary to store the full matrix *P*), but instead represent it by a low-rank approximation

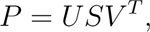

where *U ∈* R^2^*^m×r^*, *S ∈* R*^r×r^*, and *V ∈* R^2^*^d−m×r^* and *r* is the rank of the approximation. These so-called low-rank factors are stored in computer memory. If *r* can be chosen small, storing the low-rank factors results in a drastic reduction of the memory required to represent the state of the system. Since most models of signaling networks try to model the effect of an input signal on a downstream target, there is usually a couple of pathways that are described to reach this target. Even though this can also include some feedback of the pathways to the upper species, the connections within the pathways are usually from one species to the next. This results in a fairly directed graph, where most of the species have few direct connections with other species. We, therefore, predict that most signaling networks are well represented by a low-rank approximation as relatively few cuts are needed in order to partition the network. Thus, it is possible to keep the error from the partitioning low and therefore only a low rank is necessary to achieve accurate simulations.

### 2.2. Time step

Knowing the low-rank factors at one point in time we now require a way to efficiently update them according to the dynamics of the chemical master equation. To do this we employ a dynamical low-rank integrator. More specifically, the projector splitting integrator proposed in [29]. This approach is outlined in Algorithm 1.

#### Algorithm 1: Projector splitting dynamical low-rank algorithm.

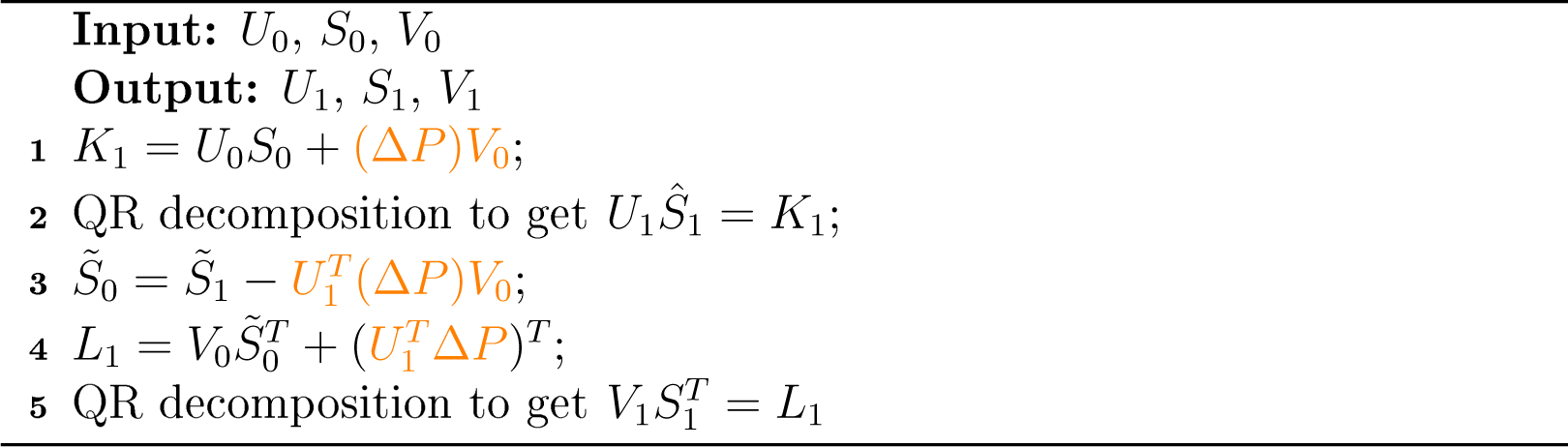

Since the chemical master equation is linear, we can write it abstractly as

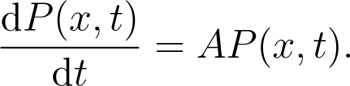

In each time step we update *P* according to Algorithm 1 where (in the simplest case of explicit Euler) we have

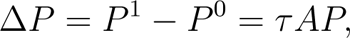

where *τ* is the time step size. The complexity of the CR decomposition is *O*(*r*^2^2*^m^*) and *O*(*r*^2^2*^d−m^*), respectively. The crucial step of the algorithm is how to compute (Δ*P*)*V*_0_*, U^T^* (Δ*P*), and *U^T^* (Δ*P*)*V*_0_ (the orange parts in the algorithm). Simply forming Δ*P* and multiplying it with *V*_0_, e.g., would be extremely expensive. That is, we have to use the structure of the chemical master equation in order to efficiently compute these three terms. This is the primary challenge that the application of the dynamical low-rank algorithm to any problem faces and we will thus address it in the following for the case of Boolean reaction networks.

### 2.3. Compute (Δ*P*)*V*_0_

We first write Δ*P* = *τ AP* = *τ A*_1_*P* + *…* + *τ A_d_P*. A rule (for more details see Appendix A) has the form *x_p_*_0_ = *F* (*x_p_*_1_ *, …, x_p__s_*), with *s* usually small and *F* a Boolean expression. Each rule can be written in matrix form (i.e. as a linear operator) as follows

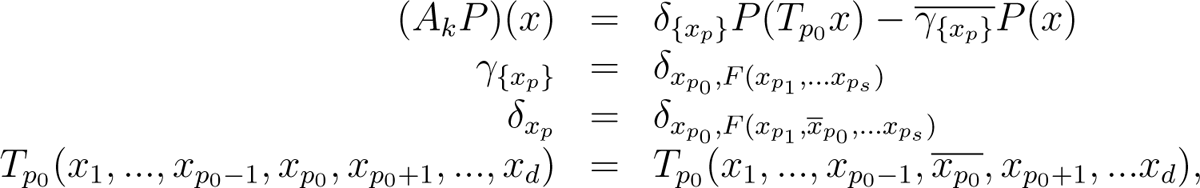

where negation is denoted by a line above the expression. We will use the convention that if *T_p_*_0_ is applied to an index that does not contain *p*_0_ it is just the identity. For simplicity, we will also write *x_G_*_1_ = (*x*_1_*, …, x_m_*), *x_G_*_2_ = (*x_m_*_+1_*, …, x_d_*), and *x_R_* = (*x_p_*_0_ *, …, x_p_*_1_).

Since there are at most *d* rules it is sufficient to consider them one by one. Then we have

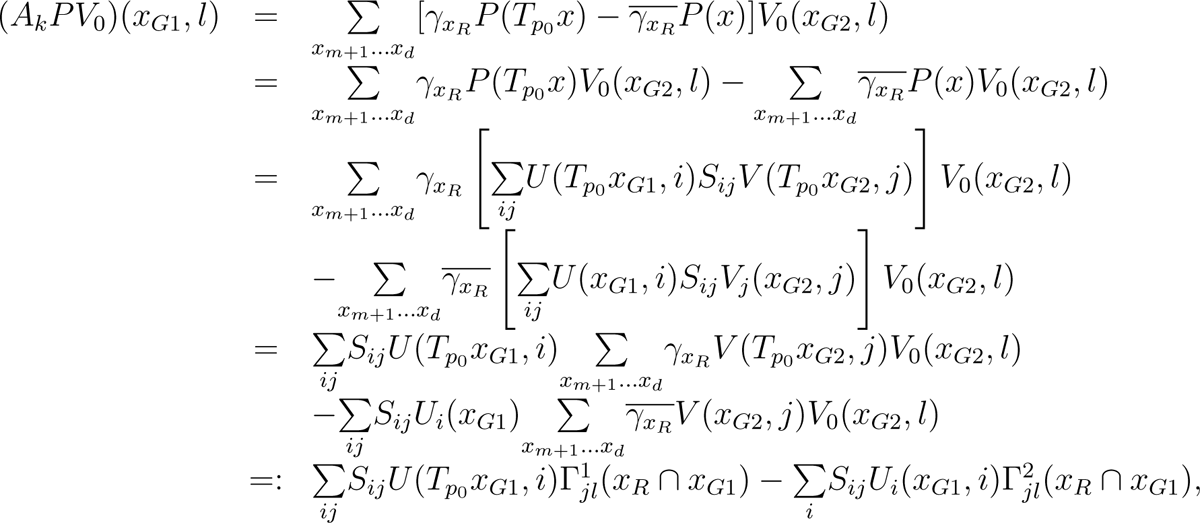

where

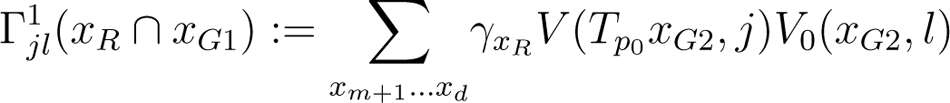

and

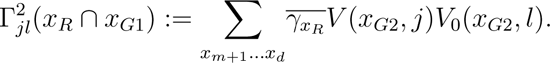

#### Algorithm 2

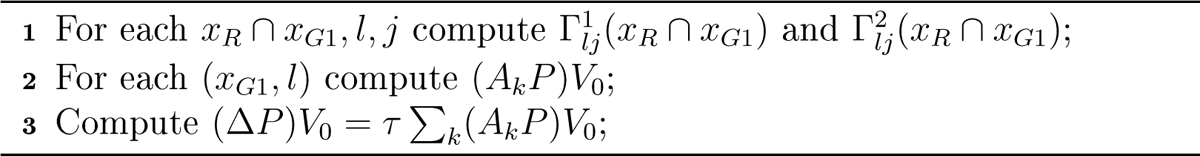

We then proceed as stated in Algorithm 2.

The first step is at most *O*(2^#(^*^xr ∩xG^*^1)^*r*^2^2*^d−m^*). The second step is at most *O*(*r*^3^2*^m^*). The third step is *O*(*k*2*^m^*). This means that the efficiency is dramatically improved as those estimates do not depend on 2*^d^*.

All of this estimates are rather pessimistic. A particular interesting case for optimization of the code is where a rule is completely contained within one partition. In this case there are two possibilities:

Case 1: *x_R_ ⊂ x_G_*_1_: We have

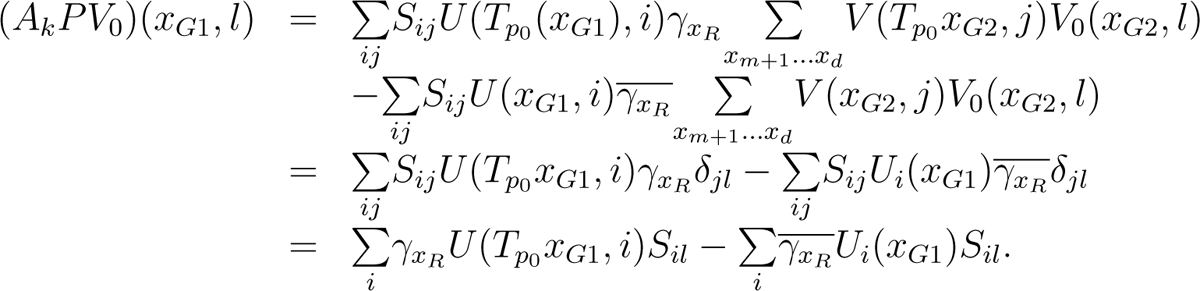

The second line follows from the orthogonality condition. This is just the direct application of the rule to *U* (i.e. no error is made). The first step in Algorithm 2 can then be omitted. The second step is only *O*(*r*^2^2*^m^*).

Case 2: *x_R_ ⊂ x_G_*_2_: In this case we have *x_R_ ∩ x_G_*_1_ = *∅* and thus the first step is only *O*(*r*^2*d*^*^−m^*).

### 2.4. Compute *U^T^* Δ*P*

We have

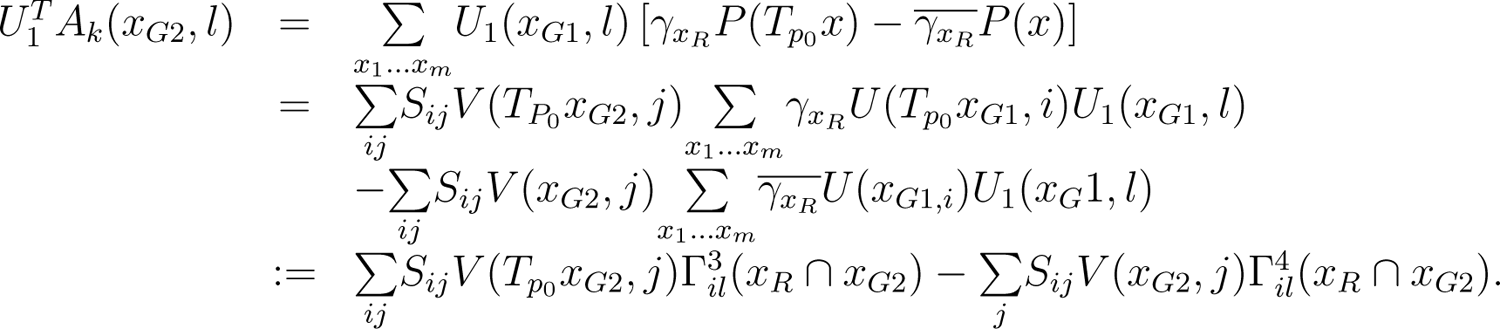

The algorithm proceeds exactly as in the previous case and we can once again make significant simplifications for the case where *x_R_* is contained within a partition. For *x_R_ ⊂ x_G_*_2_ we have

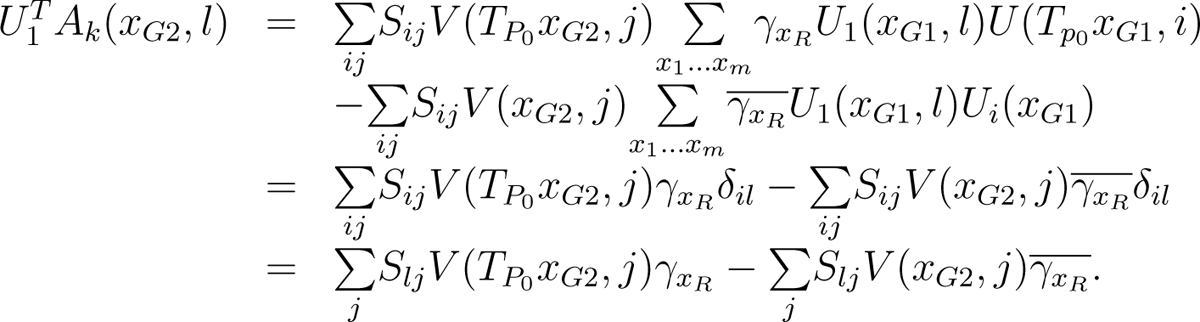

### 2.5. Compute *U^T^* Δ*PV*_0_

This is now easy since we already have computed Δ*PV*_0_ (in Subsection 2.3). So this is just a matrix-matrix product.

Before we proceed, let us make a few remarks.

Remark. The algorithm becomes more efficient if as many rules as possible are contained in one partition. This is also advantageous from an accuracy standpoint, since rules contained in one partition do not incur any low-rank error.

Remark. The notation for writing down the algorithm is somewhat complicated. However, the interpretation is very easy: if a rule is contained in one partition just apply the rule directly in the low-rank format. If not, then the states in the partition not containing *x_p_*are averaged, and the rule is applied with respect to that averaged state.

## 3. An illustrative example: a Boolean model of the mTOR pathway

To illustrate the use and efficiency of the low-rank approximation we consider a simple example; namely, the mTOR pathway model introduced by Benso et al. 6. The model includes 22 species and its network diagram is given in Figure 1. The full Boolean ruleset defining the network is as follows

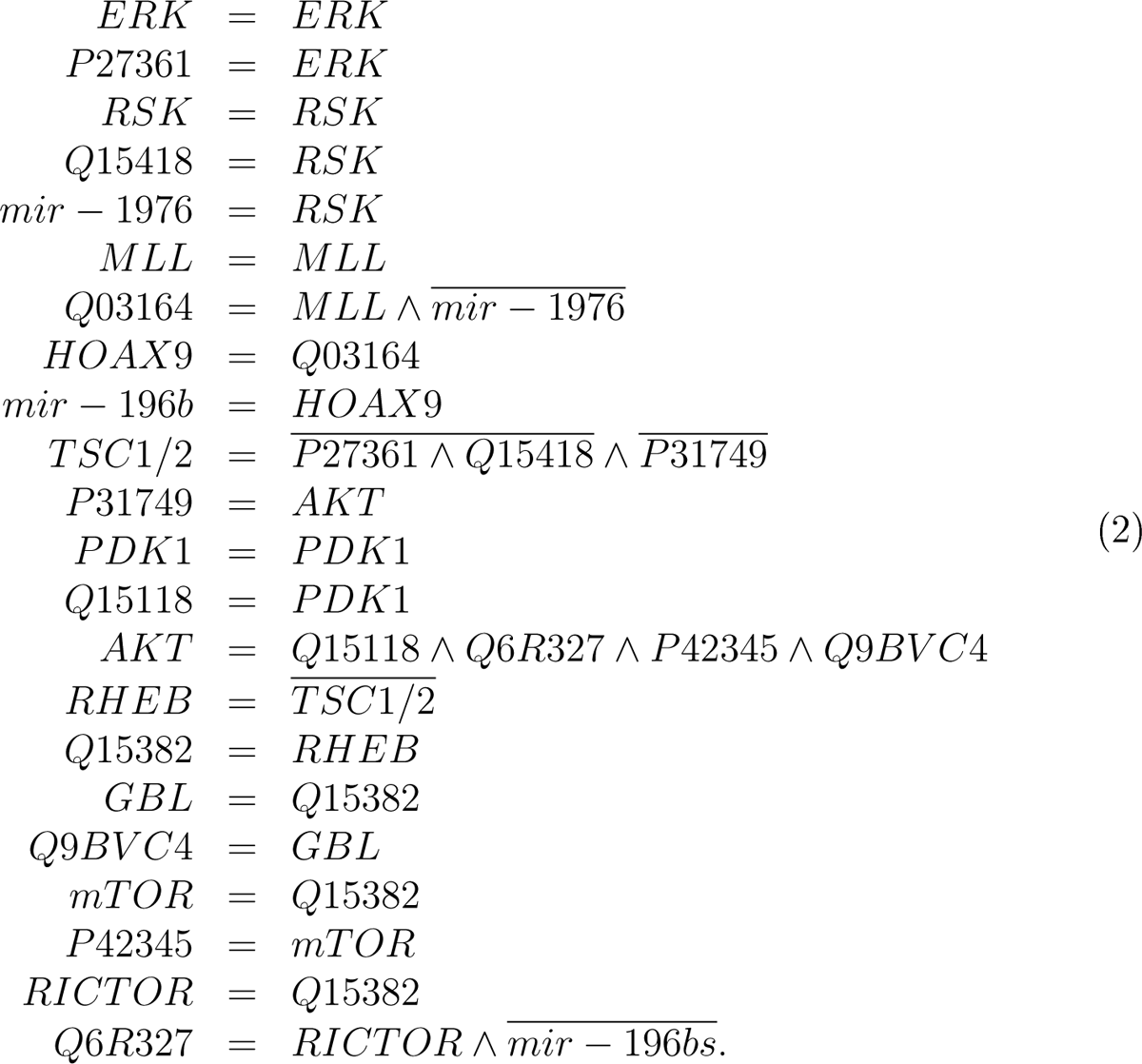

**Figure 1:**
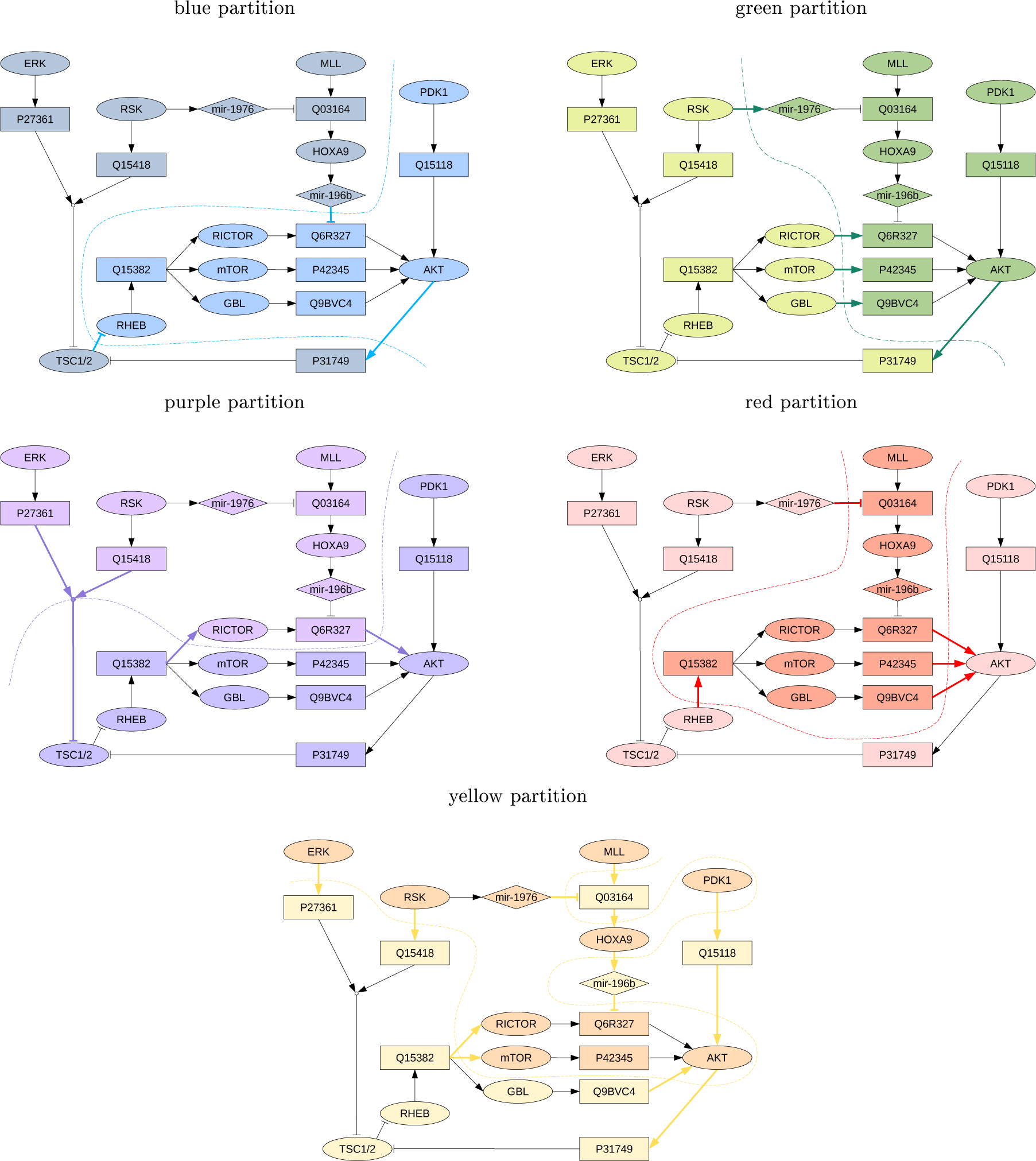
Boolean networks for the mTOR pathway model. We consider five different partitionings for the low-rank algorithms (blue, green, purple, red, and yellow). How the partitioning is done influences how small the rank (which is proportional to computational cost) can be chosen for a given accuracy. The blue and purple partitioning perform best as they treat the important pathway from Ql5382 to AKT with minimal error.

### 3.1. Network partitioning

The first step in our dynamical low-rank approach requires the partitioning of the network into subnetworks while leveraging the network structure. For this step, we require that the partitioning of the network yields subnetworks with an approximately equal number of species. Every interaction that remains within its partition (e.g. the rule for TSC1/2 in the blue partitioning in Figure 1) is treated exactly by the algorithm. Therefore, we want to minimize cross-partition dependencies (indicated by arrows in the figures) as these interactions will be subject to low-rank approximations (e.g. the pathway that connects TSC1/2 to RHEE in the blue partitioning). In Figure 1, we introduce five different partitionings that we will use in the subsequent analysis. The blue, green, purple, and red networks are chosen following this general principle of minimizing connections between the subnetworks. To demonstrate the consequences of a bad partitioning, we also included the yellow partition, where the species are randomly assigned to a partition.

The most important parameter of the low-rank approximation is the rank. A smaller rank *r* leads to faster computation times and less memory requirements for the simulation. However, decreasing rank can also lead to larger error. The maximum rank of a simulation is given by 2*^d/^*^2^, where *d* is the number of chemical species. In our example, d=22 and thus the maximum possible rank would be 2048. At the highest rank, the simulation is exact but this does not result in any computational improvements in compute time or memory use relative to a direct solver. In Figure 2, we compare the errors of five different network partitions in relation to the exact solution for different ranks (namely, *r* = 2, *r* = 4, *r* = 8, and *r* = 16). It is assumed that initially all states of the system are equally likely. For the blue (reference partition scheme) we can see that in order to obtain an error on the order of 1% (0.01) only rank 4 is required. To reduce this error to 10*^−^*^3^ the rank has to be increased to approximately 16. We also clearly see from Figure 2, by comparing e.g. the blue with the yellow partitioning, that choosing an appropriate partitioning can significantly reduce the rank and thus computational effort. Since we chose the yellow network randomly, we do not expect the partitioning to be competitive. In fact, as we can see, even increasing the rank to 16 does not decrease the error below its initial threshold, while the other partitionings see a significant reduction in the error. We also note that minimizing the connections between two partitions is not enough information in order to obtain an ideal partitioning. This can be seen in the behavior of the red and the green network. Both networks have five edges that separate the two partitions, but their accuracy is very different. The reason for this is that both the green and the red network disturb the important pathway from Q15382 to AKT in a significant way. In this case insight about the problem under consideration can help to choose an optimal partitioning and thus reduce the numerical effort.

**Figure 2:**
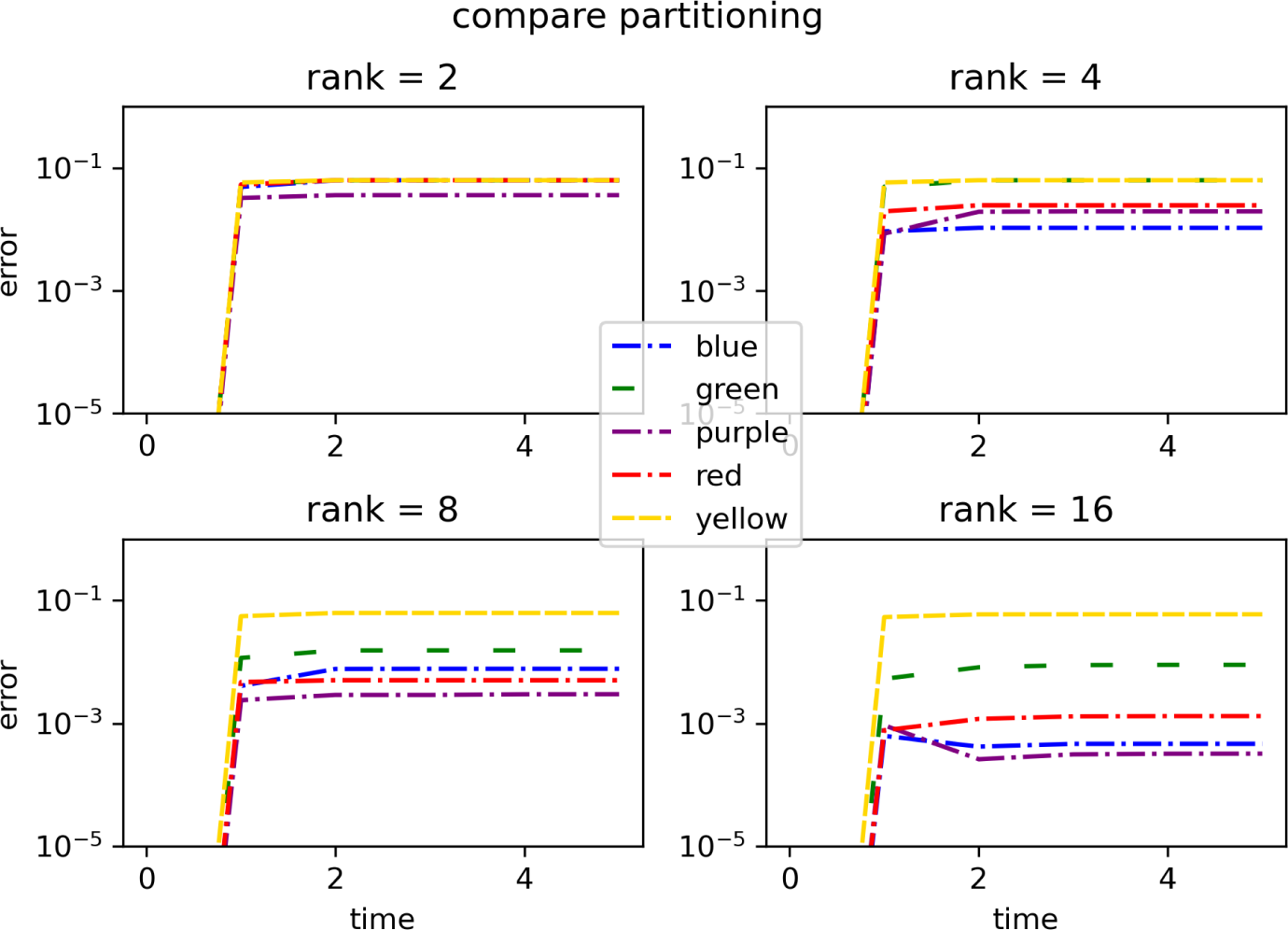
The error of the dynamical low-rank approximation for the five different partitionings of the mTOR network is shown for ranks 2, 4, 8, and 16. The error is the maximum of the difference in probability between the low-rank approximation and the exact solution at a given time (i.e. the error in *P* measured in the *L*^2^ norm).

### 3.2. Numerical results for the steady states and dynamics of mTOR

The proposed dynamical low-rank integrator is able to obtain all the statistical information in a single simulation because it is directly solving the chemical master equation. The results of the corresponding simulation are shown in in Figure 3. We display only the steady states that have a probability that exceeds 10*^−^*^3^ (20 in total for this model). For very small ranks the approximation becomes inaccurate and SS9 and SS10 are lost.

**Figure 3:**
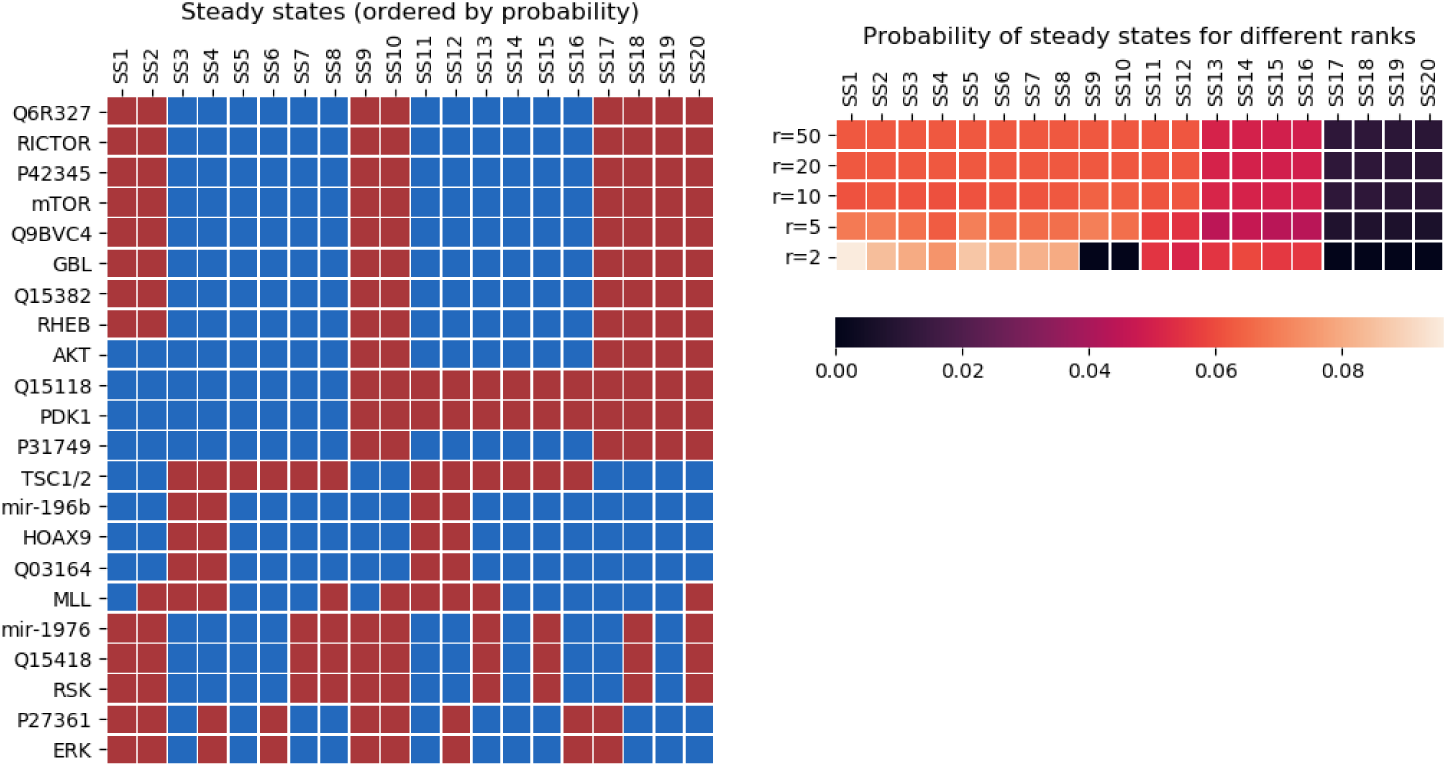
The steady states with probability larger than 10*^−^*^3^ are shown, sorted by their respective probability (red indicates on, blue indicates off). The blue partitioning and an initial value in which all states are equally likely are used.

However, even for *r* = 5 all of the relevant steady states are captured and starting at rank *r* = 10 the probabilities match exactly those of the reference solution.

In addition to the steady states, the dynamical low-rank algorithm also provides information about which trajectories lead to a given steady states as well as their probability. In order to study such time-dependent effects the dynamical low-rank approach is particularly useful. E.g. Consider the dynamics in Figure 4, where initially only a few states have a non-zero probability. Moreover, after some time the system settles in a small number of steady states. Both of these would be well represented by finite state projection. However, in the dynamics that the system follows to reach these few steady states we see an explosion in the number of states with non-zero probability. The dynamical low-rank approach, in contrast to finite state projection (which here again would suffer from the curse of dimensionality), is able to resolve these intermediate states without any increase in computational cost as we can see from Figure 4.

**Figure 4:**
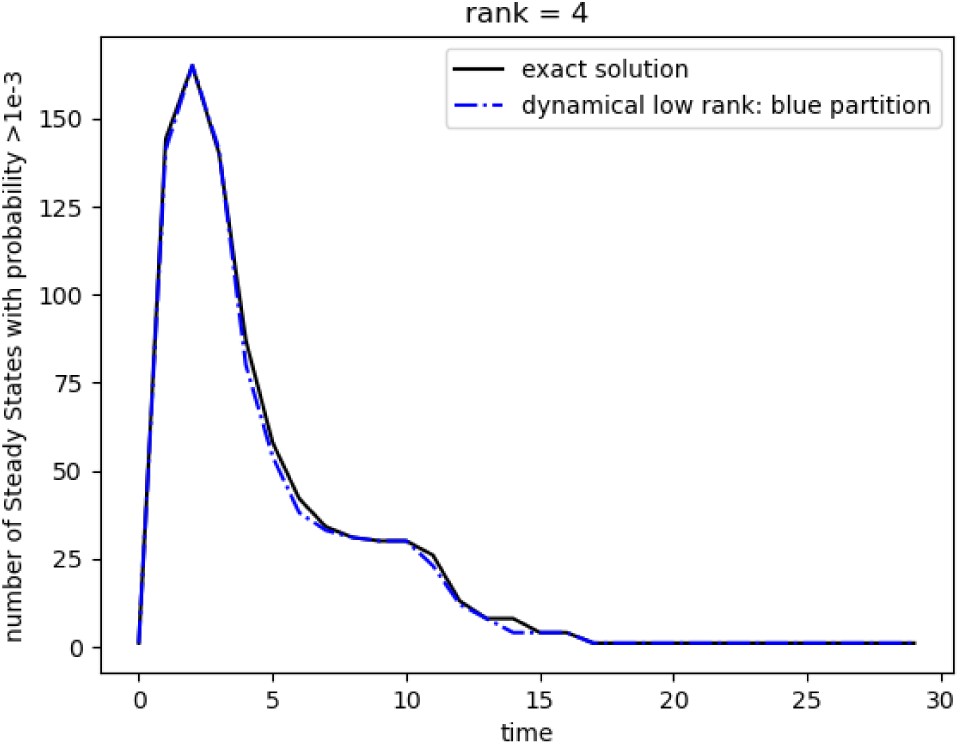
The number of states with probability larger than 10*^−^*^3^ for the mTOR network is shown as a function of time. Initially we start with a deterministic initial value (which is rank 1) and ultimately we end up in a single steady state with high probability (again rank 1). However, a large number of trajectories is generated intermittently in this dynamics.

Finally, we look at the dynamics predicted by the dynamical low-rank algorithm. In Figure 5 we plot the time evolution of mTOR for the different partition schemes and different ranks for the uniform initial condition (left) and the single initial condition (right). We observe that the blue and purple partitions in particular give results in agreement with the exact solution, even for rank 4. For rank 16 all partition schemes display the correct behavior.

**Figure 5:**
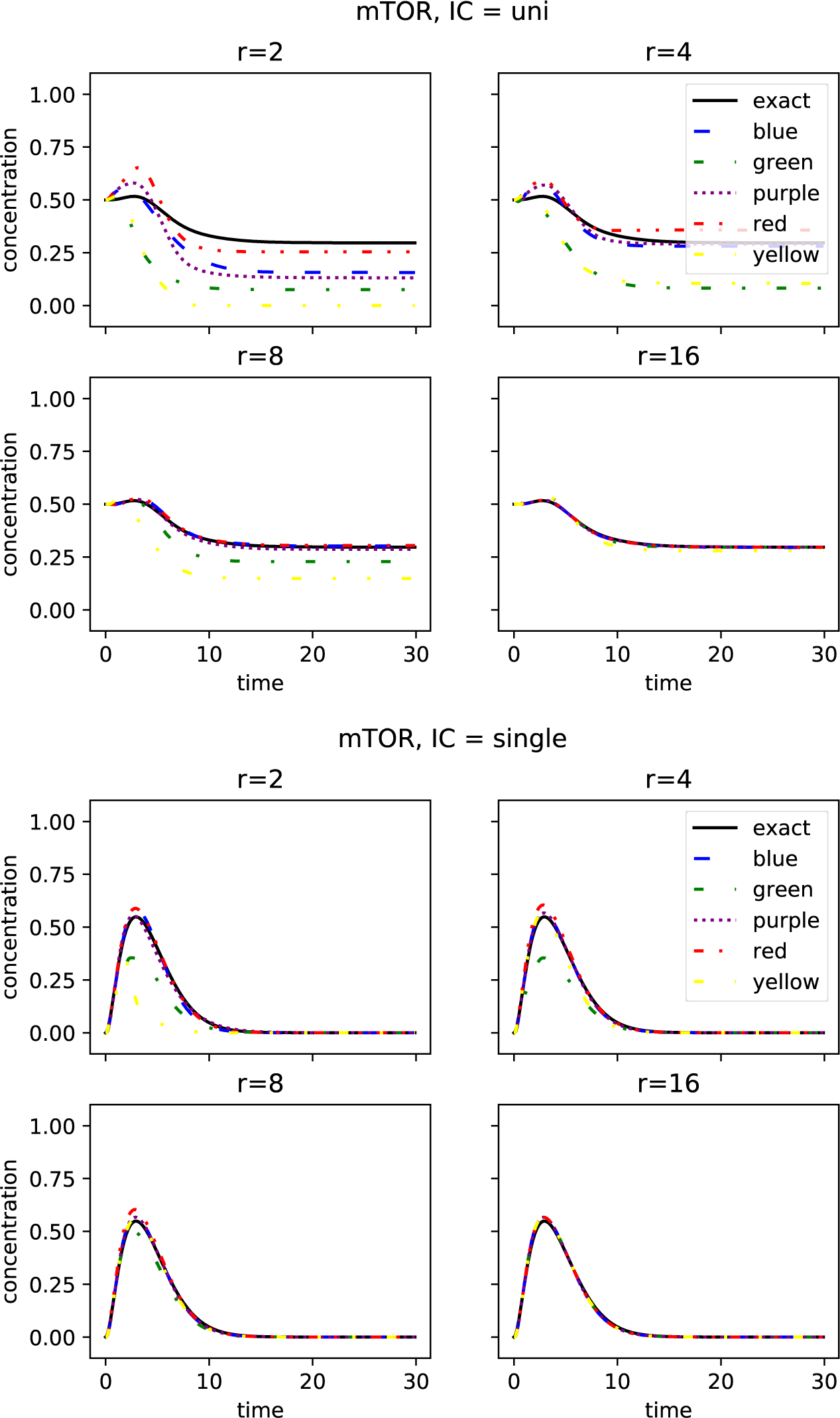
The time evolution of the probability that mTOR is switched on is shown. The configuration on the top starts with an initial configuration where mTOR initially is equally likely to be in an on or an off state. The configuration on the bottom has mTOR initially switched off and after a tra^1^n^5^sient phase that is mediated by a significant concentration of mTOR results in a steady state where mTOR is switched off again (with high probability).

We conclude that for appropriately chosen partitionings even a small rank (i.e. low computational cost) is sufficient to obtain accurate results (both in terms of steady states and dynamics). If, for some reason, a good partitioning is not available, the rank needs to be increased which adds to the computational and memory cost.

In our code, we do not only provide the projector-splitting integrator, but also more recently developed integrators, i.e., the BUG-integrator 10, as well as the augmented BUG-integrator 9. In Table 1, we compare the different performances of the integrators for solving the CME. As we can see from the table, the error of the BUG integrator is quite high. However, once we perform the augmentation, the error is almost identical to the projector splitting approach. We also note that both the BUG integrator (due to the need to recompute the coefficients for the S step) as well as the augmented BUG integrator (since the rank for the S step is increased) are somewhat more expensive compared to the projector splitting integrator.

**Table 1:**
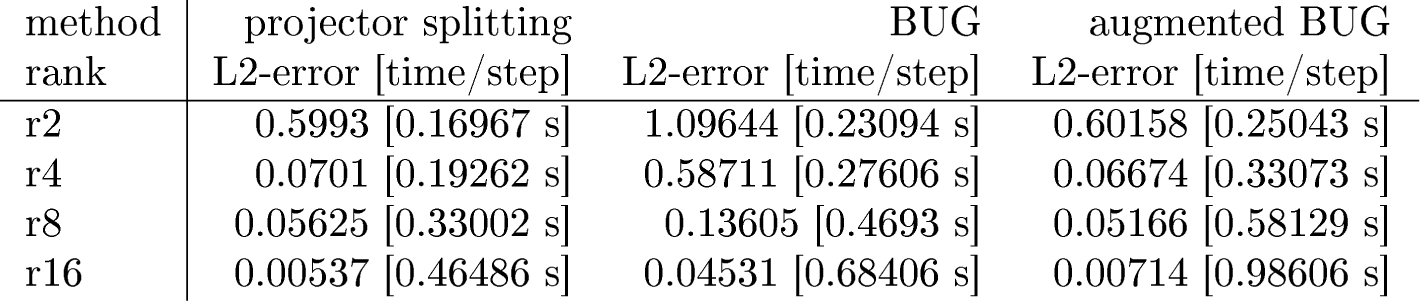
Comparison of the performance of the three different integrators that we provide in our implementation. The error is measured in the *L*^2^ norm at time point 20.

| method | projector splitting | BUG | augmented BUG |
| --- | --- | --- | --- |
| rank | L2-error [time/step] | L2-error [time/step] | L2-error [time/step] |
| r2 | 0.5993 [0.16967 s] | 1.09644 [0.23094 s] | 0.60158 [0.25043 s] |
| r4 | 0.0701 [0.19262 s] | 0.58711 [0.27606 s] | 0.06674 [0.33073 s] |
| r8 | 0.05625 [0.33002 s] | 0.13605 [0.4693 s] | 0.05166 [0.58129 s] |
| r16 | 0.00537 [0.46486 s] | 0.04531 [0.68406 s] | 0.00714 [0.98606 s] |

## 4. Computational efficiency

The mTOR network described in some detail in the previous section with its 22 species is still a relatively small model. Ultimately, we are interested in using the dynamical low-rank approach for larger problems. In the subsequent sections we will discuss two examples. A model of pancreatic cancer with 34 species and a more detailed model of apoptosis with 41 species. To illustrate the dramatic growth of memory requirements that results from increasing the number of species let us assume that a direct solver would only need to store a single input and output vector (a highly questionable assumption as a real implementation would most likely significantly exceed this theoretical value). In this case, the pancreatic cancer model would require 275 GB of main memory (RAM). This renders it beyond the scope of a desktop workstations and would require parallelization to a distributed memory supercomputer. For the apoptosis model considered the requirements would be even more severe. In fact, 35 TB of RAM would be required in this case, which is clearly not feasible.

In Table 2 (top) we show that our C--implementation of the low-rank algorithm drastically reduces these requirements. For the pancreatic cancer problem running simulations with *r* = 10, which gives very accurate results (as we will see), only 421 MB are required which is a reduction by a factor of 650. For the apoptosis model this is even more pronounced, as we would expect. In this case 12 GB are required for *r* = 10, a reduction by a factor of approximately 3000. Together with the timings provided in Table 2 (bottom) this clearly demonstrates that using the dynamical low-rank approach proposed here enables us to easily conduct these simulations on a desktop workstation within a reasonable amount of wall clock time. Thus, the only question that needs to be answered is how accurate the low-rank approximation is for realistic applications. This will be the goal of the remainder of the paper.

**Table 2:** Main memory (RAM) requirements for the mTOR pathway model (22 species), pancreatic cancer (34 species), and apoptosis (41 species) for the dynamical low-rank algorithm and different values of the rank *r* is shown on the top. For comparison, a theoretical lower bound for the memory that a direct solver would require is also provided (note, however, that an actual implementation will most likely significantly exceed this value). In all simulations the reference partitioning (blue and setup a, respectively) is used. On the bottom, timing measurements for the dynamical low-rank algorithm and different values of the rank *r* are presented. The problem is integrated until a final time *T* = 1 and all simulations have been run on a Intel Core i9-7940X CPU with 14 cores and 64 GB of DDR4 memory.

| problem | $r = 2$ | $r = 5$ | $r = 10$ | $r = 20$ | $r = 30$ | $r = 50$ | direct simulation |
| --- | --- | --- | --- | --- | --- | --- | --- |
| mTOR | 34 MB | 38 MB | 83 MB | 191 MB | – | 113 MB | > 66 MB |
| pancreatic | 160 MB | 267 MB | 421 MB | 671 MB | – | 1.4 GB | > 275 GB |
| apoptosis | 2.5 GB | 6.1 GB | 12 GB | 24 GB | 36 GB | – | > 35 TB |

Table 2: Main memory (RAM) requirements for the mTOR pathway model (22 species), pancreatic cancer (34 species), and apoptosis (41 species) for the dynamical low-rank algorithm and different values of the rank $r$ is shown on the top.
| problem | $r = 2$ | $r = 5$ | $r = 10$ | $r = 20$ | $r = 30$ | $r = 50$ |
| --- | --- | --- | --- | --- | --- | --- |
| mTOR | 0.6 s | 1.1 s | 3.4 s | 4.3 s | – | 10 s |
| pancreatic | 1 m 39 s | 2 m 55 s | 5 m 37 s | 9 m 08 s | – | 28 m 47 s |
| apoptosis | 13 m 08 s | 33 m 21 s | 1 h 06 m 33 s | 1 h 39 m 06 s | 2 h 39 m 35 s | – |

## 5. Pancreatic cancer model and some observations on asynchronous vs. synchronous updating

Results using the stochastic simulation algorithm (SSA) can be obtained using two distinct update rules. In synchronous updating all rules are applied at the same time and thus all species are updated simultaneously. While synchronous updating is commonly used, there is significant debate as to its biological interpretation [4, 20, 33, 45]. Asynchronous updating has been proposed as an alternative [39, 49, 50, 51, 53]. In asynchronous updating a single reaction is selected at random and then the corresponding rule is applied. As such, it has a direct interpretation in terms of chemical reactions and it has been shown that asynchronous updated SSA in the limit of a large number of samples gives the same dynamics as the chemical master equation [42, 47].

Nevertheless, it has been argued that synchronous updating is sufficient in order to obtain the relevant steady states [44, 43, 28, 7, 8]. However, recent work, performed using direct and SSA simulation, has called this assertion into question 21, 1. Our dynamical low-rank approach gives us a way to easily investigate the full dynamics of large systems in this context. As it is based on the chemical master equation, the dynamical low-rank approach encodes the biologically relevant asynchronous updating, while allowing us to obtain statistics over a range of trajectories and initial values with a single run of the simulation.

Gong et al. 24 use mathematical techniques in order to prove that certain properties of the Boolean network are compatible with the experimental observations. For example, it is shown that overexpression of HMGE1 will necessarily result in steady states with Proliferate switched on and Apoptosis switched off. However, these results only apply to synchronous updating. The dynamical low-rank approach, on the other hand, allows us to investigate this for the chemical master equation (i.e. with asynchronous updating). The time evolution of the probability that the cell is in the Proliferate state or that Apoptosis was triggered is shown in Figure 6. Note that we conclude from these results that in the asynchronous case the dynamics are significantly more complicated. It is possible, that apoptosis is triggered if HMGE1 is turned on (the steady-state probability is approximately 16%). Moreover, there is no difference in the ultimate cell fate (i.e. with respect to proliferation or apoptosis) whether HMGE1 is switched on or off. Thus, there is a significant difference between the steady states in the asynchronous case compared to the synchronous case that is discussed in Gong et al. 24. This casts significant doubt that the model (at least in the asynchronous case) can describe the experimentally observed effects accurately. This also highlights that models derived in the synchronous case can not (at least not without alteration) be considered a viable description of the relevant biology. Our results show that even if one is only interested in the steady states, the result for synchronous and asynchronous updating may exhibit significant differences. The dynamical low-rank approach is a tool particularly well suited for obtaining steady states that are consistent with the biologically more relevant asynchronous updating. The most likely steady states are shown in Figure 6. We once again highlight the fact that even a small rank (e.g. *r* = 5) is sufficient to capture the steady states as well as their probability very well.

**Figure 6:**
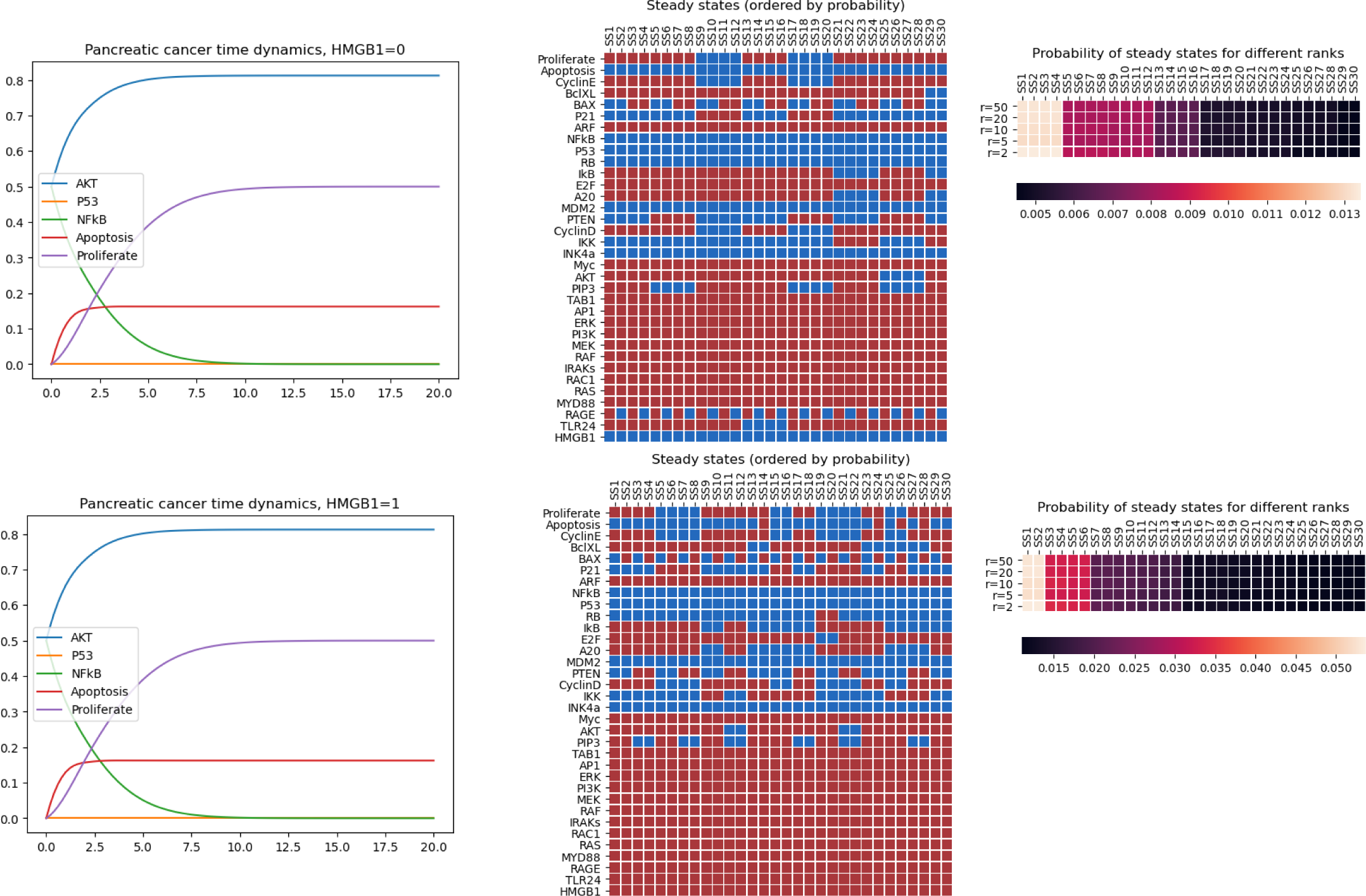
The dynamics of selected species for the pancreatic cancer model (left) and the steady sates (right) are shown for HMGBl switched off (top) and HMGB1 switched on (bottom). Steady states are sorted according to their probability. Red represents that the species is on, blue represents that the species in this steady state is off. Note, that the dynamics for Akt, p53, NFkB, Apoptosis, and Proliferate are exactly the same. The number of steady states, however, are much reduced for HMGB=l (the first couple of significant steady states have a much higher probability than for HMGB=O).

## 6. A model for cell death regulation

To demonstrate the use of a low-rank approximation in another realworld problem, we now consider a Boolean network describing programmed cell death regulation taken from Mai et al. [32]. In this model, competing signals for death and growth and their interactions are modeled to explore the mechanisms that ultimately determine the fate of the cell. The model inputs are either a signal representing Tumor Necrosis Factor (TNF) or a growth factor (GF) meant to capture stimulus by a ligand such as Epidermal Growth Factor. The main observation in the original paper, is that setting GF to “on” is ineffective in reducing apoptosis in isolation, but can reduce apoptosis execution when combined with a TNF signal. This behavior can also be observed in the simulation results presented in Figure 7.

**Figure 7:**
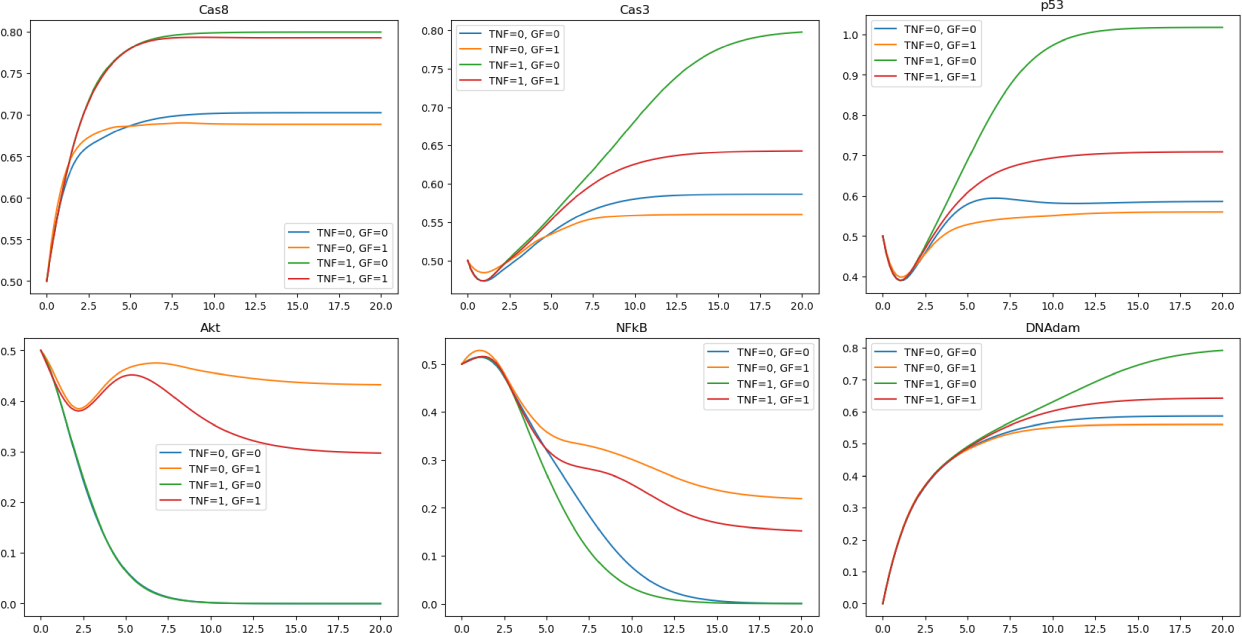
The dynamics of selected species for the apoptosis model is shown for the four different configurations of TNF and GF (i.e. TNF, GF set to 00, 01, 10, and 11).

As shown in Table 2, direct evaluation of the chemical master equation with traditional methods would require 35 Terabytes of RAM and immense amounts of computer time. However, dynamic low-rank approximation of the problem with various ranks converts the problem from intractable to easily accessible with modest computation resources on a modern desktop or laptop computer.

Figure 7 shows the dynamics of DNAdam (which in this model is a proxy for programmed cell death) alongside five of the important proteins/genes. With TNF switched off the probability of Apoptosis is approximately 60%. This is independent whether GF is on or off. Switching TNF on increases the likelihood of apoptosis dramatically. In this setting switching on GF significantly suppresses apoptosis. This is consistent with previous results from the literature 32. The mechanism of the network can be understood from Figure 8 as follows: TNF directly affects a cascade that includes various caspases that via Cas3 as the last link in the chain lead to Apoptosis. If this cascade is active AKT (which is downstream of GF) can act as an inhibitor to Cas9 breaking the direct link between TNF and Cas3. Thus, if the caspase cascade is active GF is able to suppress it. It has, however, no significant effect on the baseline Apoptosis rate (which is mediated via p53) and does not rely on the caspase cascade.

**Figure 8:**
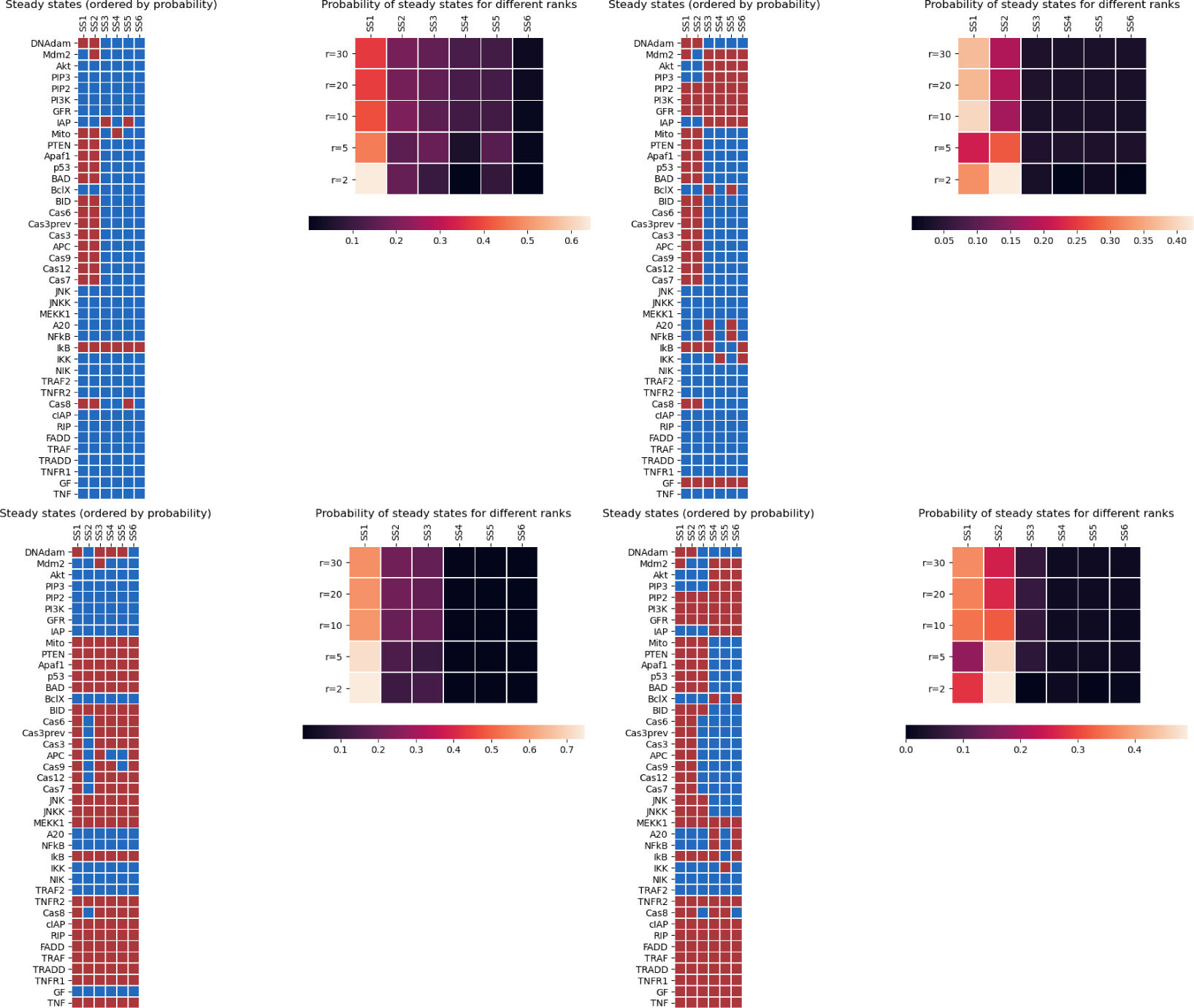
The steady states are depicted corresponding to the four different configurations of TNF and GF (i.e. TNF, GF set to 00, 01, 10, and 11; from top-left to bottom-right). The steady states are sorted according to their probability. Red represents that the species is on, blue represents that the species in this steady state is off.

## 7. Conclusion

We have shown that our dynamical low-rank approach based on partitioning the network into biological relevant parts (as opposed to use single species basis function) can achieve accurate results for the CME both in terms of dynamics and steady state results. Since a small rank (usually between *r* = 5 and *r* = 15) is sufficient, this approach results in a drastic reduction in memory and consequently allows us to solve large problems (such as the apoptosis model with 41 species for which a direct simulation would clearly be prohibitive) on a typical desktop workstation. We have also used our method to gain insight into the nature of synchronous vs. asynchronous updating, which is still heavily debated in the biochemistry community. Our work shows that for the studied pancreatic cancer model neither the steady states nor the dynamics under synchronous updating reflects well the observations made by solving the chemical master equation (i.e. asynchronous updating).

In order to enable the efficient simulation of even larger models with hundreds of species our goal is to use hierarchical tensor based dynamical low-rank approximations (such as the ones considered, e.g., in [30, 18, 31, 11]) in future work. Also in this case our viewpoint is that, using domain specific expertise, an appropriate hierarchy of subnetworks has to be identified in order to obtain accurate approximation at low rank (and thus at low computational cost).

## Code availability & validation

The code used to run the simulation and generate the plots is available at https://bitbucket.org/mprugger/low_rank_cme.

We have compared the results obtained with the dynamical low-rank implementation with a direct simulation method for a range of smaller networks (including the mTOR pathway model considered in this paper). In addition, as part of the code, a number of automated unit tests are available.

## Acknowledgments

This work was supported by National Science Foundation (NSF) CAREER award MCB 1942255 to CFL, National Institutes of Health (NIH) award 1U01CA215845 to CFL (M-PI).

## Appendix A. Translate a Boolean rule into a System of ODEs

Let us first take a look into how a system of Boolean rules can be translated into a matrix-vector multiplication. For this demonstrative purpose, we consider the small system of three species *A*, *B*, and *C* that are updated according to the following ruleset Figures A.9, A.10, A.11 demonstrate the corresponding truth tables, as well as their translation into a matrix that multiplied by the vector *P* = (000, 001, 010, 011, 100, 101, 110, 111)*^T^* recreate the rules as seen above. To update *P* according to all the rules, the matrices for each rule are added up and we get the following system of ODEs

Which is the formulation of the CME for this particular ruleset. Note, that each column/row of this matrix corresponds to a particular state of each species, e.g, the state *A* = 0, *B* = 1, *C* = 1 corresponds to the fourth column/row of the matrix *A*, (011). We therefore can see, that for a number of *d* species in the system, the formulation of the CME becomes a matrix of 2*^d^ ×* 2*^d^* entries, and thus suffers from the curse of dimensionality.

## Appendix B. Detailed error analysis

We now perform a more detailed error analysis of the dynamical low-rank approximation for different ranks and initial values. We will limit ourselves to the blue partitioning and the initial values used are summarized in Table B.3. Since the mTOR network is still small enough, we can use a reference solution obtained with a direct simulation algorithm (i.e. without performing a low-rank approximation).

The results of the comparison are illustrated in Figure B.12. For all initial values chosen the obtained results further highlight the efficiency of the low-rank approximation as in all cases a small rank is sufficient in order to obtain accurate results. The spike in the single initial condition can be attributed to the large number of intermediate states that have to be resolved at that point in time.

Figure B.13 depicts the state explosion that renders finite state projection infeasible for Boolean networks.

In Table B.4, we display the stability of the method by the mass conservation for the steady states for our five different partitions. Since the behavior of the mass is similar to the error regarding the partitioning, we consider it a good measure for the performance of partitions: Ranks 10 to 50 can not really distinguish between the blue, purple and red partition. However, while blue and red behave similar for the lower ranks, purple looses much more mass in the matter. The green partition is the least stable regarding the partitions that have not been drawn randomly, similar to Figure 2, in the main paper, where it displays the largest error for the high ranks. Unsurprisingly, the random partition performs the worst in already having a significant loss of mass for the highest rank *r* = 50 and loosing the stability with the rank completely.

## Appendix C. Partitionings for the pancreatic cancer network and the apoptosis network

### Appendix C.1. Network partitioning for the pancreatic cancer network

The pancreatic cancer network is taken from 24 and consists of 34 species. This network is used to investigate the effect of asynchronous updating versus synchronous updating. The distribution of the species we chose is shown in Table C.5 and Figure C.14.

### Appendix C.2. Network partitioning for the apoptosis networ

We chose the Boolean network with 41 nodes describing Apoptosis taken from 32. As pointed out, we would need at least 35 Terabyte of memory to compute the exact solution. Note that for our reference solution, we did not have enough memory on our system to simulate rank 50. We therefore settled for a reference solution of rank *r* = 30. Since we wanted to demonstrate how to deal with an odd number of species in our case, we also did not include the Apoptosis event, which is turned on, if the DNA Damage Event kept on in 20 continuous steps. We display the partitioning we used for our simulations in Table C.6.

**Figure A.9:**
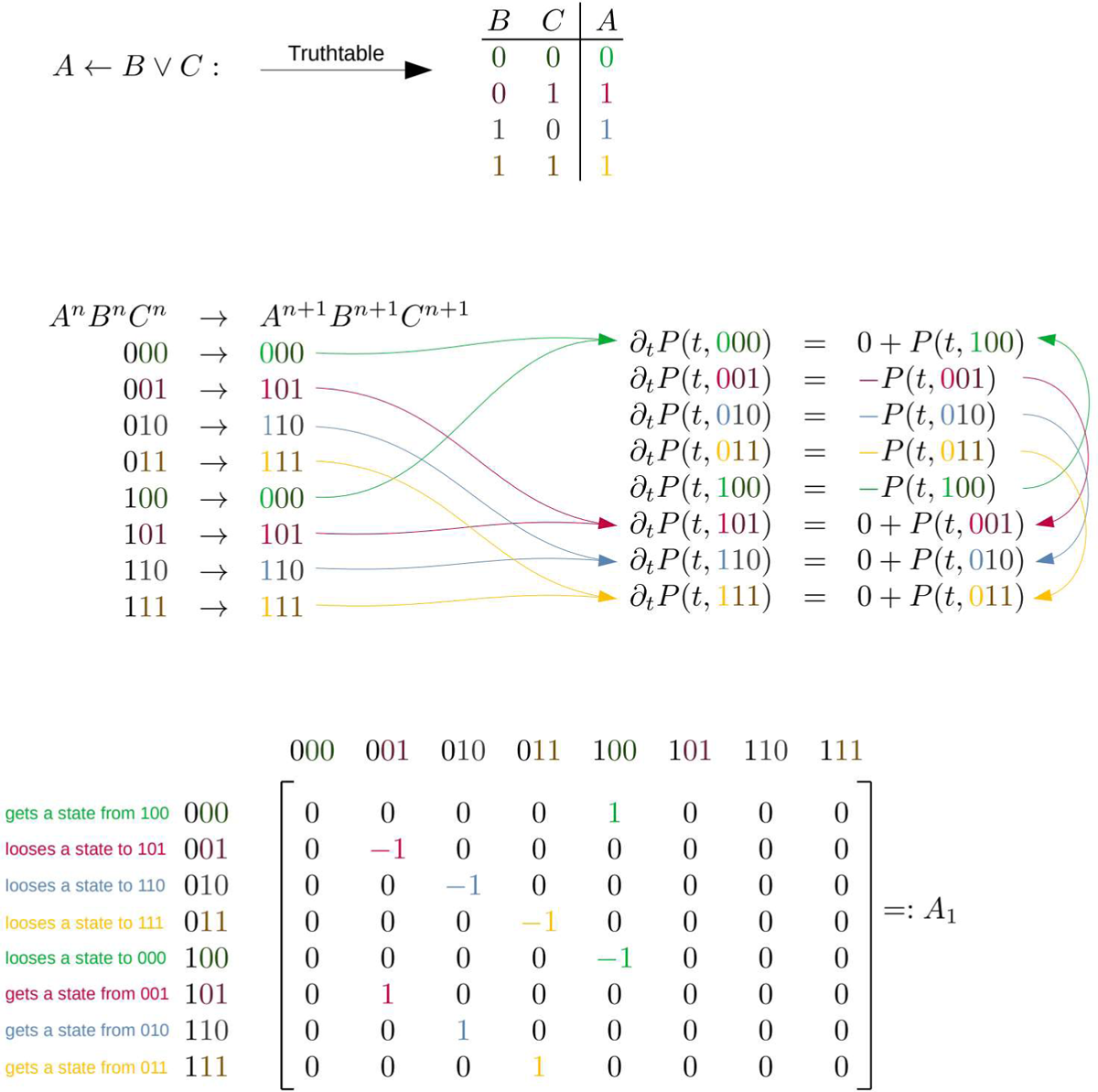
Depiction of the formulation of the rule *A ← B ∨ C* into a matrix. Note, that the negative state change from, e.g., *∂_t_P* (*t,* 001) = *−P* (*t,* 001) assures that *∂_tx_ P* (*T, x*) = 0 and thus the probability density function always stays normalized (conservation of mass).

**Figure A.10:**
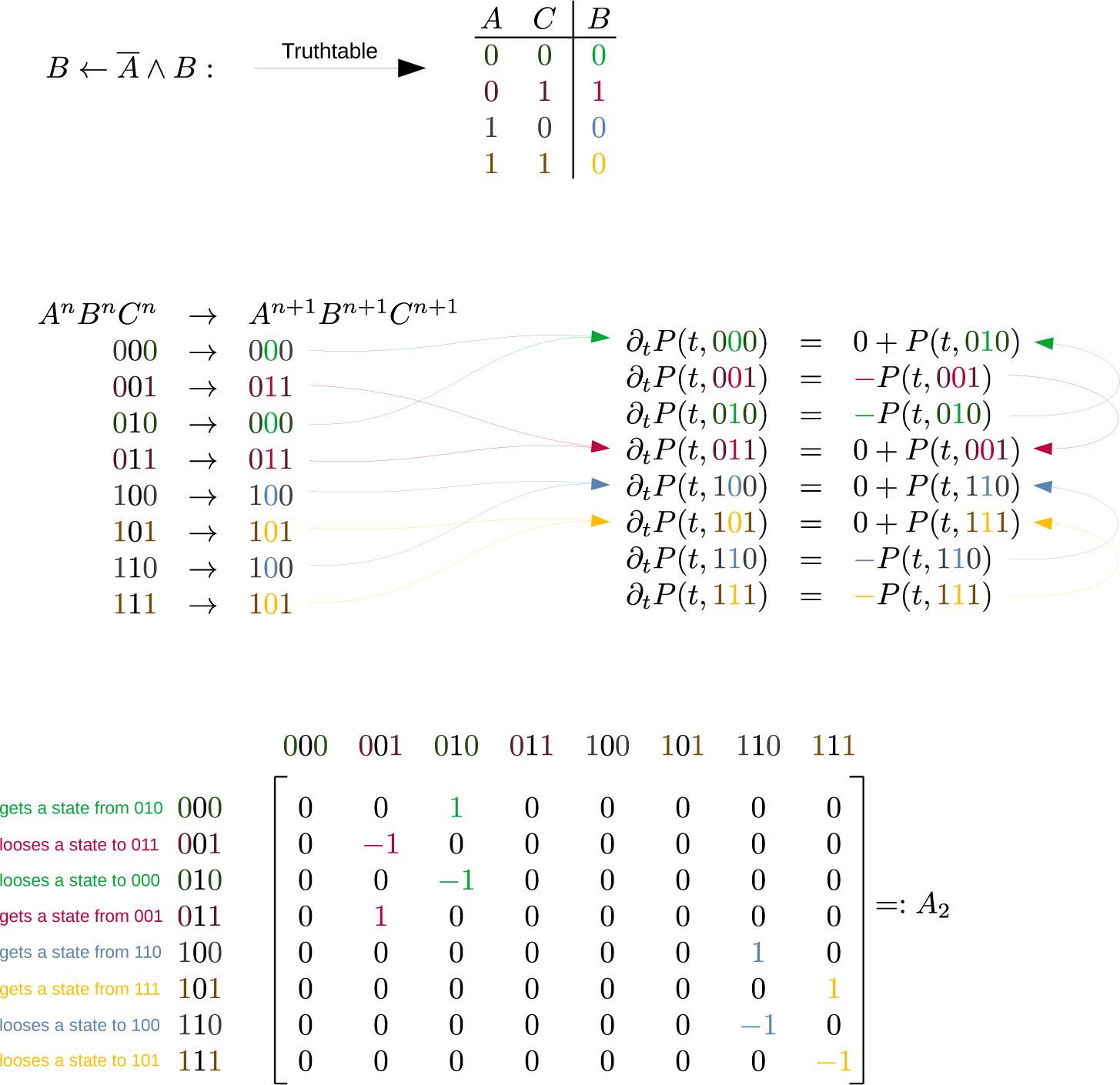
Depiction of the formulation of the rule *B ← A ∧ B* into a matrix.

**Figure A.11:**
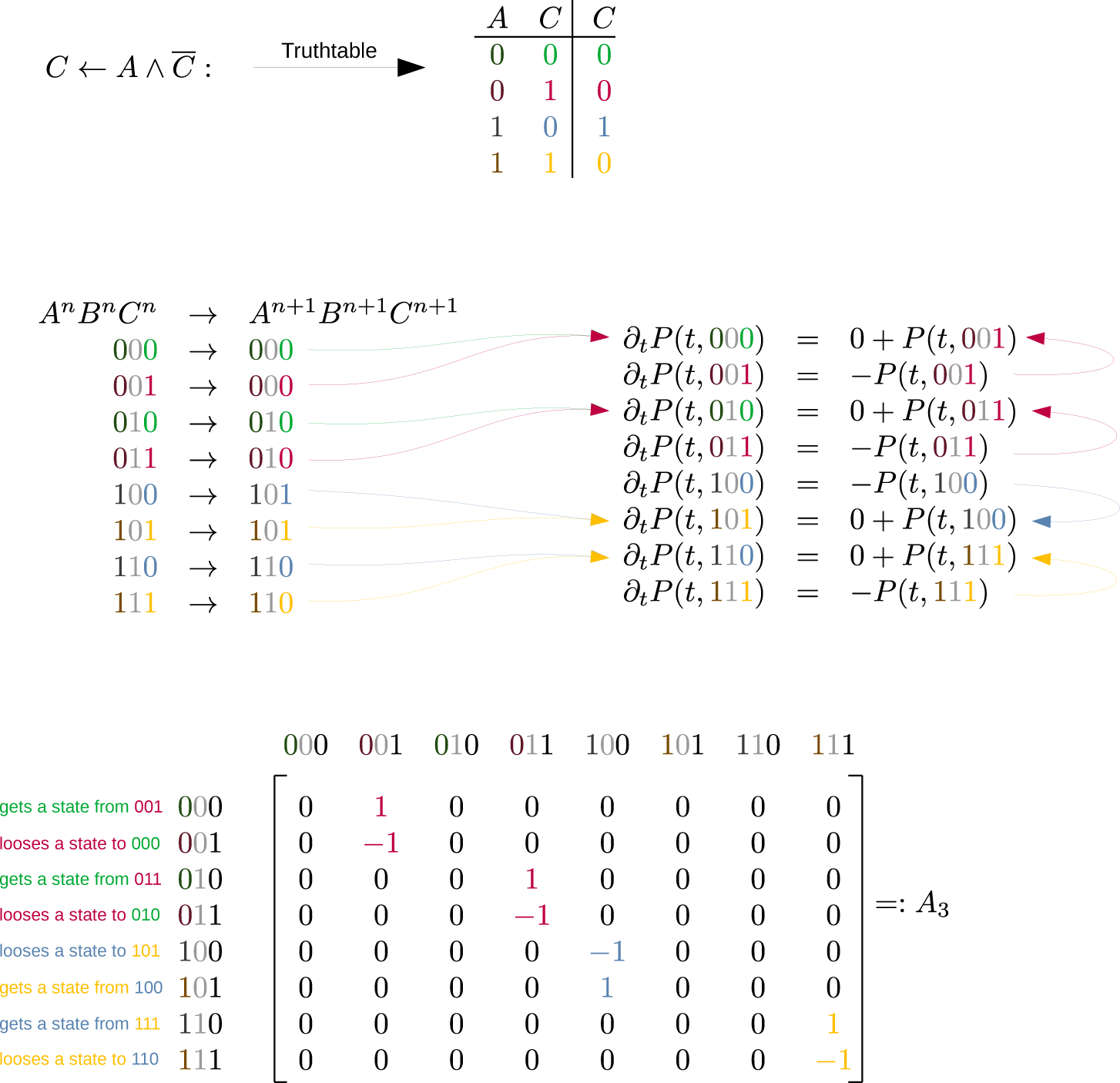
Depiction of the formulation of the rule *C ← A ∧ C* into a matrix.

**Figure B.12:**
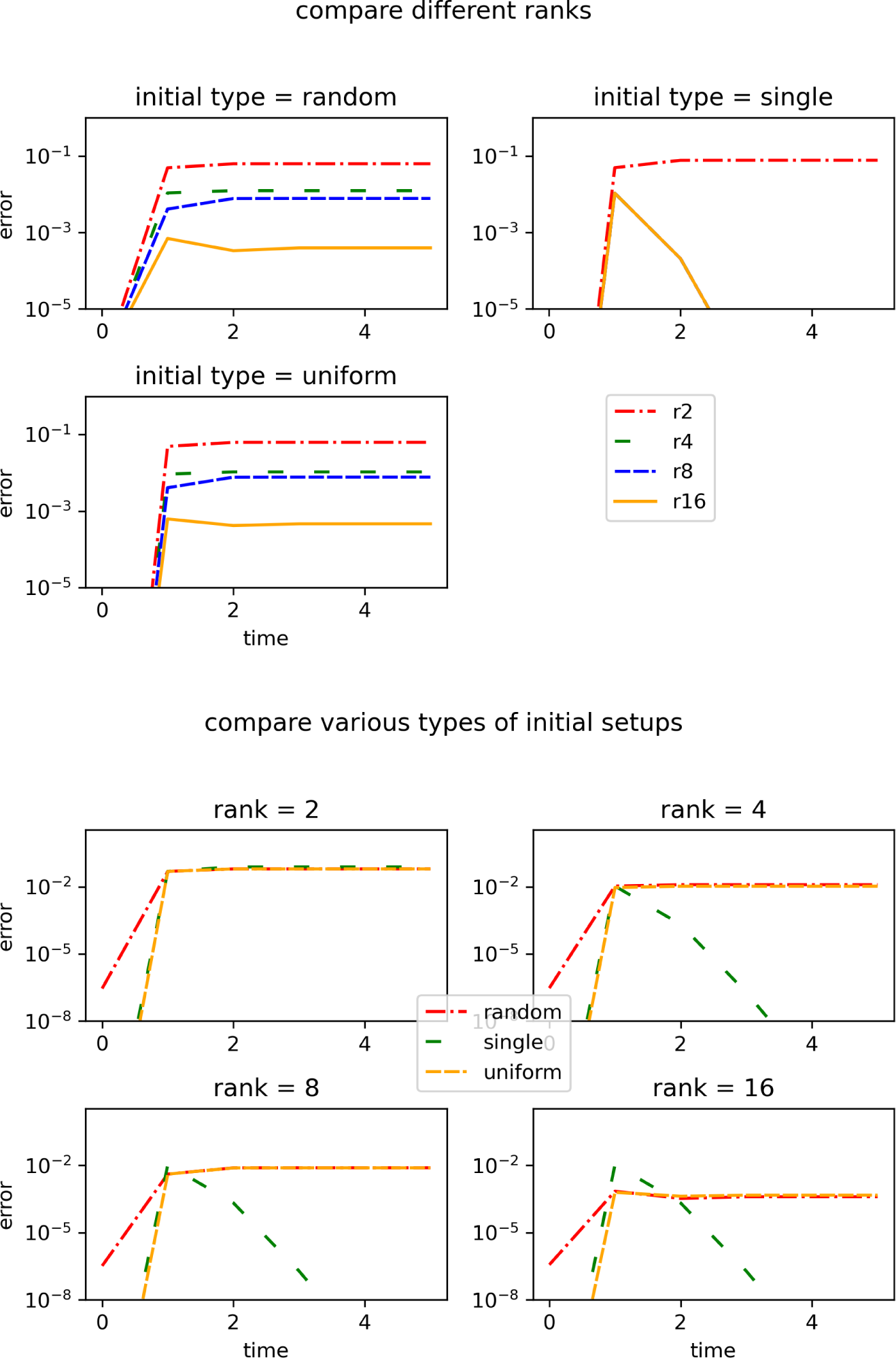
Comparison of the different setups to the exact solution according to rank and initial conditions. For the single initial setup, we picked the 387th initial vector.

**Figure B.13:**
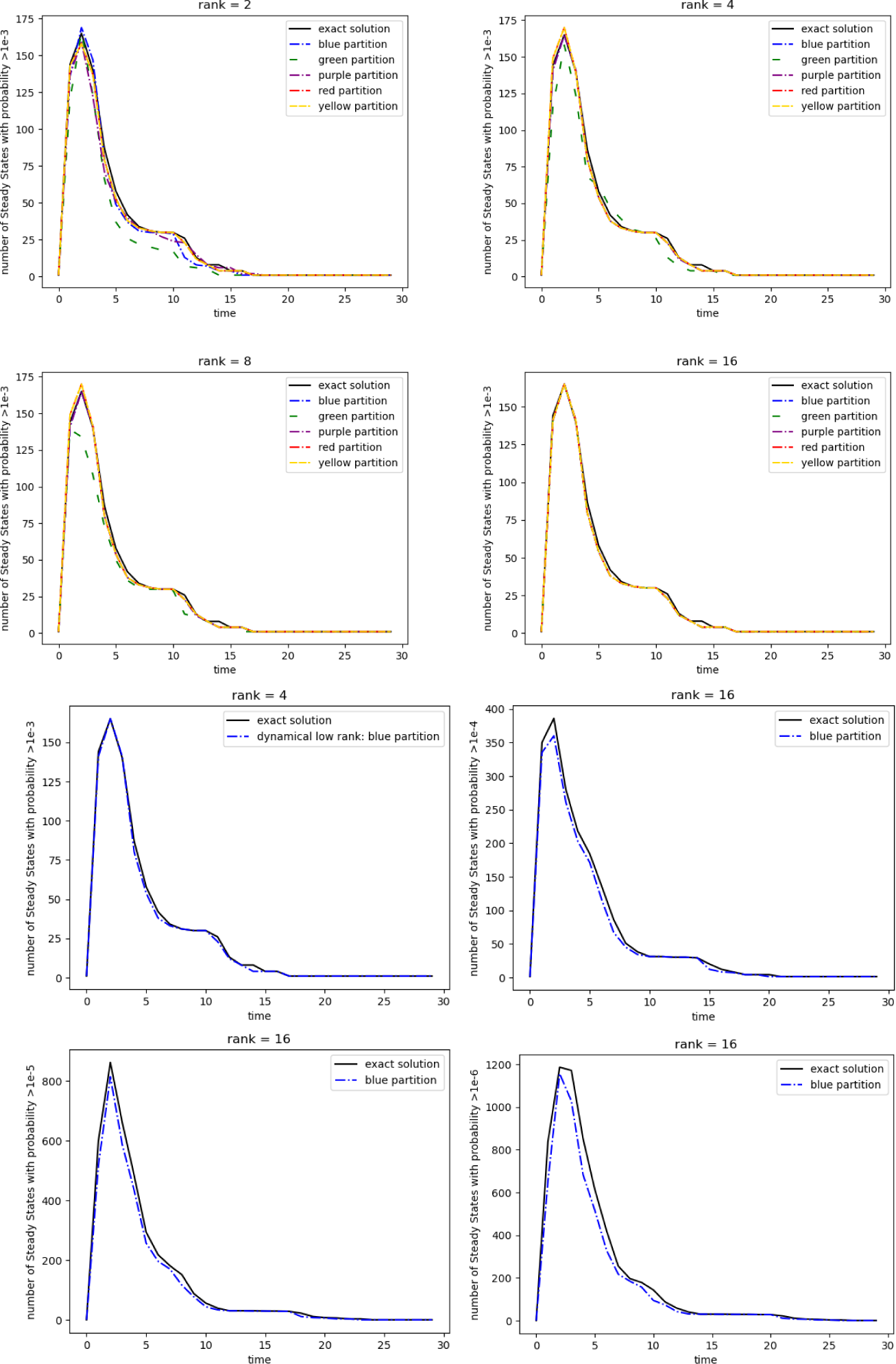
State explosion at the beginning of the mTOR network simulation for the single initial condition setup. On top, the different ranks are resolved according to the different ranks. On the bottom, for the blue partition, and rank 16, the sensitivity is lowered.

**Figure C.14:**
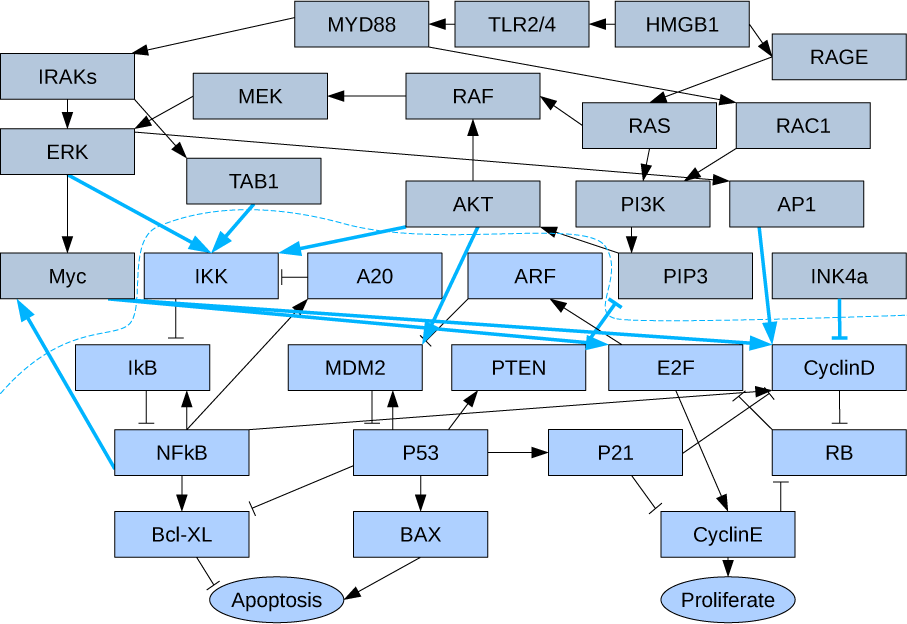
Network partitionings for pancreatic cancer taken from 24].

**Table B.3:** Initial conditions used in the analysis of the *mTOR* network.

|  |  |  |
| --- | --- | --- |
| $n = 22$ | | |
| $\text{randvec} = \text{np.random.rand}(2**n);$ | | |
| $\text{randvec} = \text{randvec}/\text{np.sum}(\text{randvec})$ | random initial setup | |
| $\text{randvec} = \text{np.zeros}(2**n);$ | | |
| $\text{randvec}[\text{randint}(0, 2**n-1)]=1.0$ | single initial setup | |
| $\text{randvec} = \text{np.zeros}(2**n);$ | single initial setup | |
| $\text{randvec}[387]=1.0$ | for comparison purposes | |
| $\text{randvec} = \text{np.ones}(2**n)/(2**n)$ | uniform initial setup | |

**Table B.4:** Steady state mass for each simulation for the network with 22 nodes.

|  | r2 | r5 | r10 | r20 | r50 |
| --- | --- | --- | --- | --- | --- |
| blue | 0.73 | 0.93 | 1.02 | 1.00 | 1.00 |
| green | 0.23 | 0.41 | 0.84 | 0.97 | 1.00 |
| purple | 0.52 | 0.85 | 0.98 | 1.00 | 1.00 |
| red | 0.70 | 0.96 | 0.99 | 1.00 | 1.00 |
| yellow | 0.10 | 0.23 | 0.27 | 0.68 | 0.89 |

**Table C.5:** Distribution of the 34 species.

|  | partition 1 | partition 2 |
| --- | --- | --- |
| species | MYD88 | IKK |
|  | TLR2/4 | A20 |
|  | HMGB1 | ARF |
|  | RAGE | IkB |
|  | IRAKS | MDM2 |
|  | MEK | PTEN |
|  | RAF | E2F |
|  | RAS | CyclinD |
|  | RAC1 | NFkB |
|  | ERK | P53 |
|  | TAB1 | P21 |
|  | AKT | RB |
|  | PI3K | Bcl-XL |
|  | AP1 | BAX |
|  | Myc | CyclinE |
|  | PIP3 | Apoptosis |
|  | INK4a | Proliferate |
| partition interactions | 10 |  |

**Table C.6:** Distribution of the 41 species.

|  | partition 1 | partition 2 |
| --- | --- | --- |
| species | TNF | Cas12 |
|  | GF | Cas9 |
|  | TNFR1 | APC |
|  | TRADD | Cas3 |
|  | TRAF | Cas3prev |
|  | FADD | Cas6 |
|  | RIP | BID |
|  | cIAP | BclX |
|  | Cas8 | BAD |
|  | TNFR2 | p53 |
|  | TRAF2 | Apaf1 |
|  | NIK | PTEN |
|  | IKK | Mito |
|  | IkB | IAP |
|  | NFkB | GFR |
|  | A20 | PI3K |
|  | MEKK1 | PIP2 |
|  | JNKK | PIP3 |
|  | JNK | Akt |
|  | Cas7 | Mdm2 |
|  |  | DNAdam |
| partition interactions | 14 |  |

## Notes

### Competing Interest Statement

The authors have declared no competing interest.

### Summary of Updates

The manuscript has been updated with timings for the unconventional low rank integrator as well as the augmented low rank integrator.

